# PARG inhibition sequesters nuclear PAR-binding proteins, including XRCC1 and its partners, into nuclear condensates to elicit cytotoxicity

**DOI:** 10.64898/2026.03.18.712393

**Authors:** Isaac Dumoulin, Brian J. Lee, Chuhan Zhang, Xiaohui Lin, Yunyue Wang, Shan Zha

## Abstract

DNA breaks activate PARP1/2 to synthesize poly(ADP-ribose) (PAR), which relaxes chromatin and recruits DNA repair factors. Normally, PAR is short-lived, rapidly degraded by poly(ADP-ribose) glycohydrolase (PARG). While PARP1/2 inhibitors are established therapies for homologous recombination (HR)-deficient cancers, predictive biomarkers for PARG inhibition (PARGi) remain undefined. Using parallel genome-wide CRISPR screens with PARP and PARG inhibitors, we show that PARGi is synthetically lethal with loss of several PAR-binding factors, including XRCC1–LIG3, POLB, ALC1/CHD1L, ARH3, and PARG itself, but notably not with HR deficiency. Conversely, loss of PARP1, NMNAT1 (required for nuclear NAD⁺ synthesis), or UNG (upstream of APE1 cleavage and PARP1 activation), confers PARGi resistance. Mechanistically, PARGi induces time- and dose-dependent formation of PARP1-and PAR-dependent nuclear condensates containing XRCC1 and associated repair factors in otherwise undamaged cells. These condensates do not harbor active DNA breaks but instead sequester PAR-binding repair proteins, depleting their available nuclear pool and impairing their recruitment to genuine DNA breaks. While our analysis focused on XRCC1, PARG inhibition likely sequesters additional PAR- and PARP1-binding proteins. Thus, we propose that PARGi sequesters PAR-binding proteins to elicit toxicity, explaining the essentiality of PARG (but not PARP1) and identifying the loss of PAR-binding factors as candidate predictive biomarkers for PARG-targeted therapy.

## Introduction

Poly(ADP-ribose) polymerase 1 (PARP1) and PARP2 are DNA damage-activated ADP-ribosyl transferases that are rapidly recruited to and activated by DNA strand breaks through direct recognition of 5′-phosphorylated DNA ends, a molecular moiety exposed upon DNA breakage^1,2^ Once activated, PARP1 and PARP2 catalyze the synthesis of poly(ADP-ribose) (PAR) on themselves and other substrates, including histones and XRCC1, using NAD⁺ as the ADP-ribose donor and releasing nicotinamide^3^. PAR, a highly negatively charged polymer with nucleic acid-like properties, can form condensates *in vivo* and *in vitro*^4^. PAR facilitates DNA repair by promoting chromatin relaxation and recruiting DNA repair factors through their PAR binding motifs, including the single-strand DNA break (SSB) repair scaffold protein XRCC1, implicated in ligation during base excision repair (BER) and beyond^5,6^. Dual PARP1 and PARP2 inhibitors (PARPi) block the enzymatic activity of PARP1/2 and delay single-strand DNA break repair. Cancer cells defective in homologous recombination (HR), most notably those lacking BRCA1-PALB2-BRCA2, are hypersensitive to PARPi, a vulnerability that was initially attributed to the conversion of unrepaired single-strand breaks into replication-associated double-strand breaks (DSBs), which rely on HR for resolution^7–9^. Beyond catalytic inhibition, clinically approved PARPi prolonged the appearance of PARP1/2 at DNA lesions (termed “trapping”)^10–12^, where they obstruct DNA repair and replication, contributing to both anti-cancer therapeutic effects and the dose-limiting toxicity of PARP inhibitors^13–16^. Over the past decade, four PARPi have been approved for the treatment of BRCA-deficient ovarian and other cancers^17^. Despite robust initial responses, resistance inevitably develops, most commonly through BRCA1/2 reversion mutations^18,19^, motivating the development of additional PAR-targeted strategies. In this context, small inhibitors for poly(ADP-ribose) glycohydrolase (PARGi) have been developed^20^.

PARG is the principal enzyme that degrades long poly(ADP-ribose) (PAR) chains^20^. The released ADP-ribose (ADPr) can be recycled into ATP and ultimately feed the NAD+ salvage pathway through NAMPT and compartmentalized nicotinamide mononucleotide adenylyltransferases, including nuclear NMNAT1^21^. PARG uses its macrodomain to engage the terminal ADPr and efficiently removes ADPr units in an exonucleolytic manner. However, it cannot efficiently hydrolyze the final mono-ADPr attached to amino acid side chains or nucleic acids; those terminal adducts are instead removed by TARG1 or ARH3^22,23^. Several small-molecule PARG inhibitors (PARGi) have since been developed^24–28^. Early models proposed that preventing PAR degradation might reduce nuclear NAD+ availability^21,29,30^, thereby indirectly limiting PARP1/2 activity in a manner analogous to PARPi, which preferentially target HR-deficient cancer^31,32^. However, several lines of evidence suggest that this model is overly simplistic. First, nuclear NAD+ serves as a critical cofactor for multiple enzymes in addition to PARP1, including the Sirtuin family of histone deacetylases. Moreover, NAD+ recycling requires only one molecule of nicotinamide released by PARP1/2 and two molecules of ATP, which can be supplied by diverse metabolic pathways^33^. Although PARG generates ADP-ribose (ADPr) that can contribute to ATP production, nuclear ATP concentrations (∼5 mM) significantly exceed NAD+ levels (100–150 μM)^34^ making it unclear whether PARG inhibition reduces ATP sufficiently to impair nuclear NAD+ regeneration. Consistent with this view, NAMPT inhibition (FK866) causes a more pronounced and sustained depletion of cellular NAD+ than PARG inhibition but does not produce comparable proliferation defects,^35^ suggesting that PARGi toxicity involves mechanisms beyond NAD+ depletion. Genome-wide CRISPR screens using PARG inhibitors or inducible PARG degradation (dTAG-PARG) further indicate that NAD+ metabolism is not the limiting determinant of PARG sensitivity^35^. Moreover, although PARP1 is dispensable for murine development, PARG knockout leads to early embryonic lethality^28,29^, highlighting essential functions beyond simply counteracting PARP1. While PARGi can target PARPi-resistant BRCA-deficient cancer cells through a replication-associated mechanism^36^, the molecular basis of this vulnerability and the fundamental mechanism underlying PARGi toxicity remain largely undefined.

To delineate the cellular impact of PARG inhibition and understand the difference between PARGi versus PARPi, we performed parallel genome wide CRISPR screens in mammalian cells^37,38^ using three published PARG inhibitors (PDD00017273, COH34, and JA2131^26–28^) alongside three FDA-approved PARP inhibitors (olaparib, niraparib, and talazoparib), all at their IC90 concentrations. As expected, loss of PARP1 conferred resistance to both PARGi and PARPi. In addition, loss of NMNAT1, which catalyzes nuclear NAD+ synthesis, also confers PARGi resistance, arguing against nuclear NAD+ as a limiting factor. In contrast to PARPi, PARGi did not exhibit synthetic lethality with HR deficiency. Instead, the strongest synthetic lethal interactions with the best characterized PARGi, PDD00017273, involved loss of XRCC1 and its partners LIG3 and POLB, as well as ALC1, all PAR-associated factors. Mechanistically, PARG inhibition induced the formation of PARP1- and PAR-dependent nuclear condensates containing PARP1, XRCC1, POLB, LIG3, and PARP2 in otherwise undamaged cells. These condensates were dynamic but lacked PCNA, suggesting that they may represent remnants of DNA damage foci following repair completion. With prolonged PARG inhibition, these condensates sequester nuclear PARP1 and XRCC1, impairing their recruitment to newly arising DNA lesions and thereby sensitizing cells to single-strand break-inducing agents (*e.g.*, MMS), as seen with XRCC1-deficiency^39^. Consistent with this model, gRNAs targeting UNG, PARP1, or NMNAT1 conferred resistance to PARGi, suggesting that spontaneous PARP1 activation at endogenous lesions, such as UNG-initiated nicks, drives PAR-dependent condensate formation under PARG inhibition. Together, these findings support a model in which PARG inhibition causes the formation of PARP1- and PAR-dependent nuclear condensates that sequester XRCC1–LIG3–POLB, ALC1, along with other PAR-binding proteins, after the completion of DNA repair, thereby providing a unified mechanistic explanation for the broad and severe toxicity of PARG inhibitor that is fundamentally distinct from PARP inhibition.

## Results

### Parallel genome-wide CRISPR screens reveal a distinctive vulnerability to PARG inhibition compared with PARP inhibition

To compare the impact of PARGi versus PARPi, we performed whole-genome CRISPR screens with an independently derived mouse gRNA library in v-abl kinase-transformed murine B cell line carrying the Emu-Bcl2 transgene^40^ **(Fig. 1A).** These cells were selected because they are diploid, maintain a stable karyotype, and can be efficiently infected and sorted^36,39^. Moreover, it offers the opportunity to carry out the screen in an independent murine gRNA library^37,38^. Specifically, cells containing an inducible Cas9 were transduced with a genome-wide lentiviral gRNA library targeting 18,424 mouse genes (4–6 gRNAs per gene)^38,41,42^. Cas9 expression was induced with doxycycline for 5 days, a subset of the library was sequenced to identify clones harboring guides targeting essential genes, then the rest of the pool libraries were divided and grown for 6 days with either DMSO or IC90 concentration of PARPi or PARGi **(Fig. 1A).** IC90 concentrations were determined and validated by MTT assay in parallel conditions on the same CRISPR/Cas9-expressing clones **(Fig. S1A-G).** We chose IC90 to focus on rate-limiting vulnerabilities of PARG and PARP inhibitors in short-term proliferating cell culture. Data were analyzed using the MAGeCK pipeline, which considers both the consistency and the impact of the gRNAs targeting each gene^43^. The β- and z-scores were calculated by comparing drug-treated and DMSO-treated conditions. We tested all three published PARG inhibitors available at the time - PDD00017273^26^, COH34^28^, and JA2131^27^ - together with three FDA-approved PARPi - olaparib, niraparib, and talazoparib. However, in two technical replicates, the commercially purchased COH34 exhibited inconsistent behavior **(Fig. S1H, Table S1)** and seems to target other PAR metabolic enzymes *(e.g.*, DPH family). JA2131 was a generous gift from Dr. John Tainer’s lab with limited availability **(Table S1)**. Thus, we focused on the best characterized and commercially available PDD00017273 (referred to as PDD thereafter).

**Figure 1.**
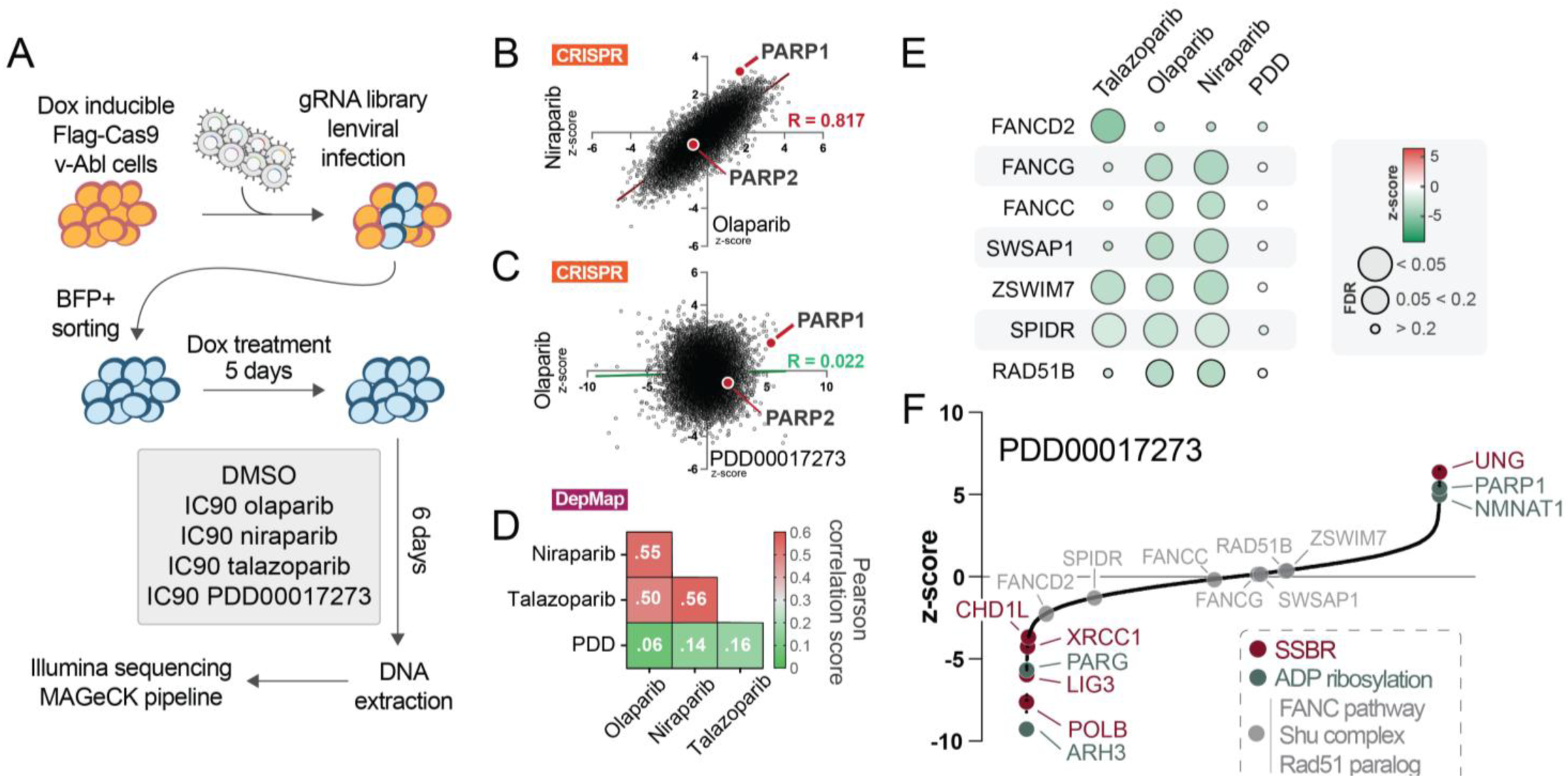
Parallel CRISPR Screen of PARP and PARG inhibitors. **A.** V-abl B cells expressing Flag-tagged Cas9 were infected with a BFP-expressing lentiviral gRNA library. BFP+ cells were sorted and treated with doxycycline for five days to induce Cas9 expression. Indicated inhibitors treatment was performed at IC90 for six days. Then, DNA was collected from the cells for sequencing and analysis using the MAGeCK pipeline. **B.** Scatter plot of total CRISPR genes Z-scores for Niraparib and Olaparib IC90 treatments. Simple linear regression and Pearson correlation test were performed to determine R. **C.** Scatter plot of total CRISPR genes Z-scores for PDD0017273 and Olaparib IC90 treatments. Simple linear regression and Pearson correlation test were performed to determine R. **D.** Summary table of Pearson R coefficient for linear regression between cell lines sensitivity to PDD, olaparib, nirabarib and talazoparib. Depmap sensitivity score (PRISM repurposing data) of 536 cell lines to the indicated compound. Simple linear regression and Pearson correlation test were performed to determine R. Colored scaled is represented on the right and R coefficients are written in white. **E.** Heatmap representing z-score (color range) and False Discovery Rate FDR (size) for the genes targeting indicated in rows, in cells challenged with inhibitor indicated above, in columns. Gene implicated in homologous recombination are displayed. **F.** CRISPR screen z-scores ranking of co-essential genes with PDD00017273 treatment. SSBR related genes are shown in red, ADP-ribosylation related genes are shown in dark blue, HR related genes are shown in light grey.

Consistent with prior literature^35,44–47^, loss of PARP1 causes resistance to both the PARPi and PARGi screens (**Fig. 1B, Table S1)**. Meanwhile, PARP2 is not a significant hit in any screens, likely reflecting its less robust activity and the fact that PARP2 recruitment to DNA damage sites, in part, depends on PARP1 activity^48^ **(Fig. 1B)**. Linear regression analysis of the CRISPR screen Z-scores for two PARPi, olaparib and niraparib, yielded a strong correlation (Pearson r = 0.817; **Fig. 1B**). In contrast, the correlation between PDD and olaparib scores was low (Pearson r = 0.022; **Fig. 1C**). Similar results were obtained by comparing PDD vs niraparib or PDD vs talazoparib (**Fig. S1I-J**). In addition, DepMap analyses of PDD and three PARPi on > 500 cancer cell lines^49,50^ also show a similarity among three PARPi (pairwise Pearson r = [0.5 - 0.56]; **Fig. 1D, S1K-L**) and poor correlation between PARPi and PDD (pairwise Pearson r = [0.06 - 0.16]; **Fig. 1D, S1H-I**). Thus, despite shared vulnerability profiles among PARPi, PARG inhibition (e.g., PDD00017273) elicits cellular responses that are mechanistically distinct from PARP inhibition.

### PARG inhibition reveals selective vulnerability of single-strand break repair, but not homologous recombination

PARPi preferentially target HR-deficient cancers, particularly those harboring BRCA1/2 loss-of-function mutations^7–9^. While BRCA genes are essential in the v-abl kinase-transformed murine B cells and not identified as significant hits in the CRISPR screens, loss of the alternative HR mechanisms - SWSAP1-ZSWIM7-SPIDR (SHU complex)^51^, RAD51B^52^ or Fanconi anemia (FA) pathway^53–55^ - all show hypersensitivity to PARPi **(Fig. 1E, S2A-C)**. Meanwhile, these HR-related genes were not significant in the PARGi/PDD screen (**Fig. 1E, S2D**), indicating that PARG inhibition is not synthetically lethal with loss of these HR factors under the same conditions. Instead, mono-ADPr hydrolase ARH3 is the top synthetic lethal target in PDD-treated cells (**Fig. 1F**), consistent with its overlapping role with PARG in ADP-ribose hydrolysis^56^ and in removing ADPr/PAR from protein substrates. Loss of PARG itself also exhibited strong synergistic lethality with PDD (**Fig. 1F**). Similar results were observed in an independent CRISPR screen performed in HEK293^35^, and have been attributed to the incomplete loss of PARG function following CRISPR-mediated disruption of this large gene, whose catalytic domain resides in the C-terminus (∼97 kb in mouse and ∼123 kb in human genomes). Notably, both PARG and ARH3 also bind to PAR, especially the terminal ADPr.

Meanwhile, gRNAs targeting multiple components of the base excision repair (BER) pathway emerged among the most prominent synthetic lethal hits. During BER, DNA glycosylases recognize damaged bases and generate abasic sites, which are processed by APEX1 to generate a nick with a 3′-OH and a 5′-deoxyribose phosphate, an intermediate that activates PARP1. In our screen, loss of the uracil DNA glycosylase (UNG), which functions upstream of APE1 cleavage and PARP1 activation, conferred strong resistance to PARGi, phenocopying PARP1 loss. No other DNA glycosylases in our libraries were significant hits for either PARGi or PARPi **(Fig. S2E)**, suggesting that uracil-associated lesions constitute a major source of spontaneous PARP1 activation in these cells. In contrast, loss of downstream BER factors, XRCC1-LIG3-POLB resulted in marked hypersensitivity to PARGi **(Fig. 1F)**. This polarized pattern of genetic interactions, with resistance conferred by disruption of UNG and PARP1, and hypersensitivity arising from loss of downstream repair factors (XRCC1-LIG3-POLB), is consistent with uracil excision-triggered PARP1 activation as a primary source of toxicity in PARGi-treated cells. Supporting this model, loss of NMNAT1 that catalyze nuclear NAD+ synthesis from NMN and ATP^57^, but not cytoplasmic NMNAT2 or mitochondrial NMNAT3, conferred resistance to PARGi/PDD **(Fig. 1F, Table S1)**, indicating that nuclear PARP1 activation, rather than global NAD+ or ATP depletion, is the critical determinant of PARGi sensitivity.

Given that PAR promotes recruitment of the POLB-XRCC1-LIG3 complex to DNA damage sites, it was initially perplexing that PARG inhibition, which leads to PAR accumulation, would be synergistically lethal with POLB-XRCC1-LIG3 loss. Notably, the loss of another PAR-binding protein, ALC1 (CHD1L), also exhibited strong synthetic lethality with PARGi **(Fig. 1F)**, raising the possibility that PARG inhibition induces a functional shortage of PAR-binding DNA repair factors. Consistent with their PAR-dependent recruitment, loss of POLB or ALC1 also sensitizes cells to PARPi **(Fig. S2F)**. However, losses of XRCC1 and LIG3 show no significant genetic interaction with PARPi^58^ **(Fig. S2F)**, despite the well-characterized role of XRCC1 in recruiting POLB^54^. In this context, prior studies demonstrated that XRCC1 deficiency leads to PARP1 hyperactivation, such that PARP1 deletion or inhibition rescues the growth defects associated with XRCC1 loss^39,59,60^, providing one explanation for the absence of XRCC1–LIG3 in PARPi screens. In addition to enzymatic inhibition, PARPi are known to block DNA replication by “trapping” both PARP1 and PARP2 at the DNA lesions, where they block DNA repair^10,11,46,61^. In this regard, PARG does not have a DNA-binding domain and is recruited to sites of DNA damage through MACRO domain-ADP-ribose (ADPr) interaction^20^, rendering direct trapping on DNA unlikely, which might provide one explanation for the absence of HR in the PARGi screen. Given the central role of PAR in recruiting POLB-XRCC1-LIG3 and ALC1 and the requirement for PARG-mediated PAR turnover to resolve PAR-dependent assemblies, we propose a model in which PARG inhibition delays PAR degradation and sequesters PAR-binding repair factors, including XRCC1, LIG3, POLB, and ALC1, within PAR-dependent nuclear condensates that persist after repair completion.

### PARGi induces XRCC1-PARP1 condensation in undamaged cells

To test the sequestration hypothesis, we expressed C-terminal GFP-tagged PARP1 (PARP1-GFP) and N-terminal RFP-tagged XRCC1 (RFP-XRCC1) in *PARP1^KO^* RPE1 cells^46^. Strikingly, even in the absence of any micro-irradiation, 10 µM PDD induced a time-dependent accumulation of PARP1 and XRCC1 coalescing into nuclear condensates, detectable within 15 minutes and peaking at ∼24 hours **(Fig. 2A-D)**. Condensation was quantified as the ratio of cumulative condensate-integrated intensity to total nuclear intensity (see Methods). Co-treatment with the dual PARP1/2 inhibitor niraparib effectively abrogated PDD-induced PARP1 and XRCC1 condensates **(Fig. 2A-B)**, indicating that PARP1/2 catalytic activity, and by extension PAR synthesis, is required for condensate formation. In contrast to micro-irradiation–induced canonical PARP1 foci, which consistently co-localize with the DNA damage marker PCNA^12^, PDD-induced PARP1 condensates lacked RFP-PCNA **(Fig. 2E-F, S3A-B)**. The PCNA foci typically persist for at least 30 minutes after micro-irradiation^46^. Thus, PDD-induced PARP1-XRCC1 condensates do not represent canonical DNA damage foci but instead arise independently of DNA damage or persist long after repair completion.

**Figure 2.**
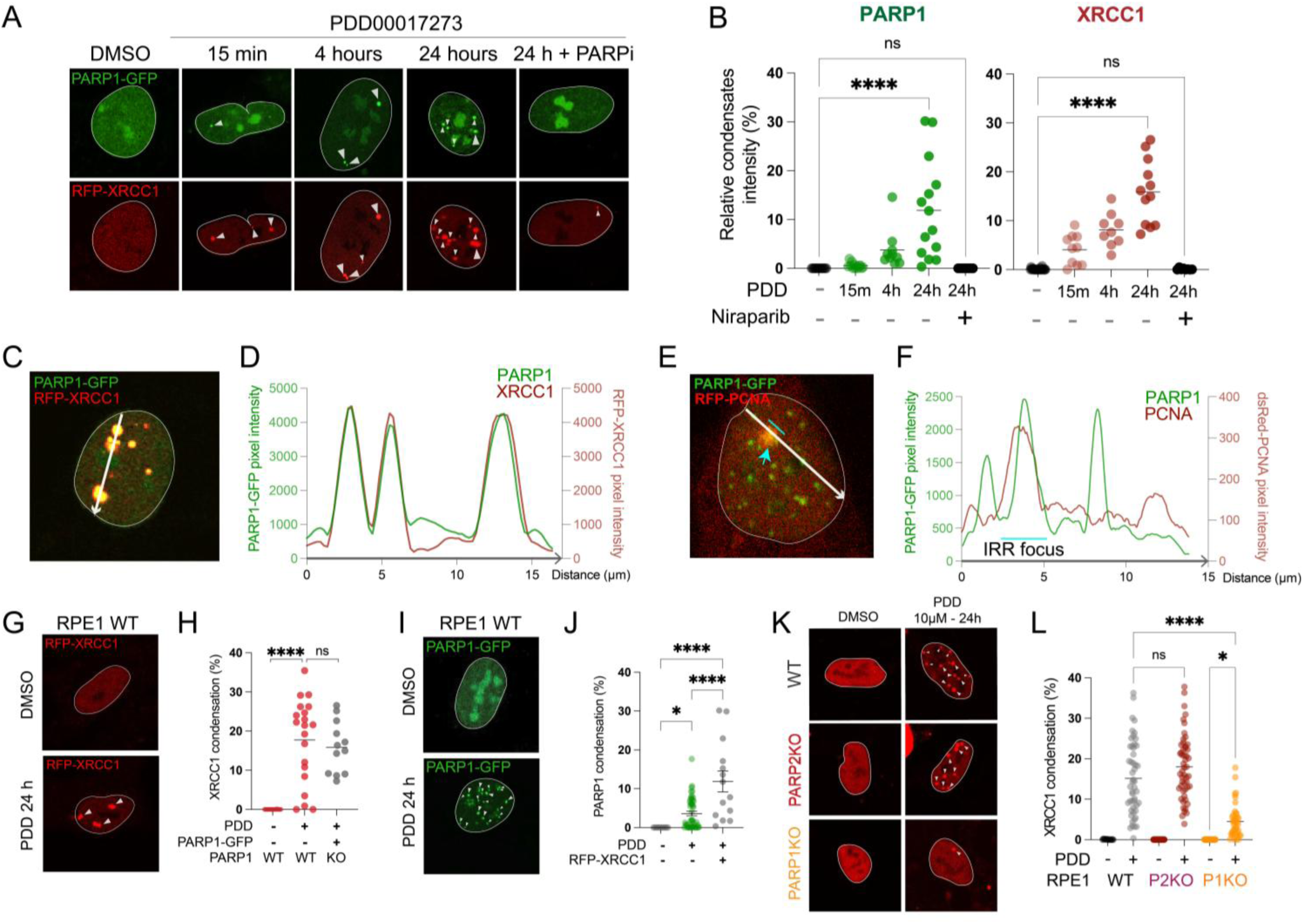
PARG inhibition causes time dependent nuclear co-condensates containing both PARP1 and XRCC1, but not PCNA. **A.** RPE1 PARP1KO cells transfected with GFP-PARP1 and RFP-XRCC1. PDD PARGi were used at 5 µM (15 minutes, 4 hours) and 10 µM (24 hours). PARPi (niraparib) was used at 1 µM for 24 hours. Nuclei are encircled with a thin white border for easier visualization and white arrowheads point at foci. **B.** Quantification of RFP-XRCC1 (red) and GFP-PARP1 (green) foci intensity over the total nuclear intensity. **C.** RPE1 PARP1KO cells transfected with PARP1-GFP and RFP-XRCC1 and treated with PARGi for 24 hours (10 µM). White arrow represents the quantification profile highlighted in D. **D.** Representation of intensity profile for PARP1 (green) and XRCC1 (red) for C. **E.** RPE1 PARP1KO cells transfected with PARP1-GFP and RFP-PCNA and treated with PARGi for 4 hours (5 µM) before micro-irradiation using 405nm laser. White arrow represents the quantification profile highlighted in D. and blue line and arrow highlights the zone of micro-irradiation. **F.** Representation of intensity profile for PARP1 (green) and PCNA (red) from E. **G.** Representative images of RFP-XRCC1 nuclear distribution in DMSO or PDD treated RPE1WT cells. White arrows point at the condensates. **H.** Representation of the condensate intensity over the total nuclear intensity of the indicated protein. Data in grey was already used in B. **I.** Representative images of PARP1-GFP nuclear distribution in DMSO or PDD treated RPE1WT cells. White arrows point at the condensates. **J.** Representation of the condensate intensity over the total nuclear intensity of the indicated protein. Data in grey was already used in B. **K.** Representative images of RFP-XRCC1 nuclear distribution in DMSO or PDD treated RPE1WT (grey), PARP1-KO (red) or PARP2-KO (orange) cells. White arrows point at the condensates. **L.** Representation of the condensate intensity over the total nuclear intensity of the indicated protein. All statistical analysis (B, H, J, L) were performed using ordinary one-way ANOVA with Šídák’s multiple comparisons test (*p ≤ 0.05, **p ≤ 0.01, ****p ≤ 0.001, ****p ≤ 0.0001)

Because phase separation can depend on protein abundance, we tested whether PDD-induced XRCC1 condensates require ectopic PARP1 expression. XRCC1 condensates formed similarly in wild-type cells and in PARP1^KO^ cells reconstituted with GFP–PARP1 **(Fig. 2G-H)**, indicating that PARP1 overexpression is not required. Consistently, PDD induced condensates of RFP-tagged murine XRCC1 in immortalized mouse embryonic fibroblasts (iMEF) (**Fig. S3C-D**), suggesting conservation across species. In the reciprocal experiments, XRCC1 overexpression enhanced PARP1 condensate formation but was also not required for it **(Fig. 2I-J)**. In *XRCC1^KO^* cells, PARP1 condensates were also more pronounced, consistent with PARP1 hyperactivation in the absence of XRCC1^39^ **(Fig. S3E-F).** Interestingly, we noticed that the number of PDD-induced PARP1 condensates seems to correlate with XRCC1 expression, with the most in XRCC1^KO^ cells, intermediate in WT cells, and the fewest upon XRCC1 overexpression **(Fig. S3E-G)**. Thus, although XRCC1 is not required for PDD-induced PARP1 condensate formation, it modulates PARP1 condensate level and size by regulating repair kinetics/PARP1 activity and beyond. Finally, we expressed RFP–XRCC1 in *PARP1^KO^* or *PARP2^KO^* RPE1 cells to assess the relative contributions of PARP1 and PARP2. Whereas PARP2 loss had minimal impact, PARP1 null reduced XRCC1 condensate formation by ∼4-fold **(Fig. 2K-L)**, indicating that PARP1 and its catalytic activity are the primary drivers of PDD-induced PARP1–XRCC1 condensation. Taken together, PARG inhibition induces prominent PAR-dependent nuclear PARP1–XRCC1 condensates independent of ongoing breaks.

### PARGi-induced XRCC1-PARP1 condensates show similar dynamics to micro-irradiation-induced PAR-foci

To determine whether PDD-induced PARP1-XRCC1 condensates are dynamic or represent irreversible aggregates, we performed fluorescence recovery after photobleaching (FRAP)^46,61^ following 24-hour PDD treatment and compared these to canonical micro-irradiation–induced (IRR) foci. PARG inhibition did not alter the recovery kinetics of free nuclear PARP1 (t_1/2_ < 5s; maximal recovery > 90%) (**Fig. 3A-C**). In contrast, PARP1 within PDD-induced condensates exhibited slower recovery (t_1/2_ ∼ 15.28 s), comparable to that observed at canonical micro-irradiation sites (t_1/2_ ∼15 s), indicating that these condensates are not irreversible aggregates but resemble damage-induced assemblies. However, maximal recovery was reduced in PDD-induced condensates (∼75% versus ∼95% at damage sites), suggesting that a fraction of PARP1 is less mobile or not readily exchangeable. In general, XRCC1 is more dynamic (t_1/2_ < 1s in nucleoplasm vs ∼4s for PARP1), but displayed a similar pattern, with slightly slower yet efficient partial recovery in the condensates (t_1/2_ ∼2.14s versus ∼0.76 s; maximal recovery 75% versus ∼100%) **(Fig. 3D-F)**. Together with the lack of PCNA, these findings support a model in which PARG inhibition drives the persistence of PARP1- and PAR-dependent nuclear condensates that represent residual DNA damage assemblies following repair completion.

**Figure 3.**
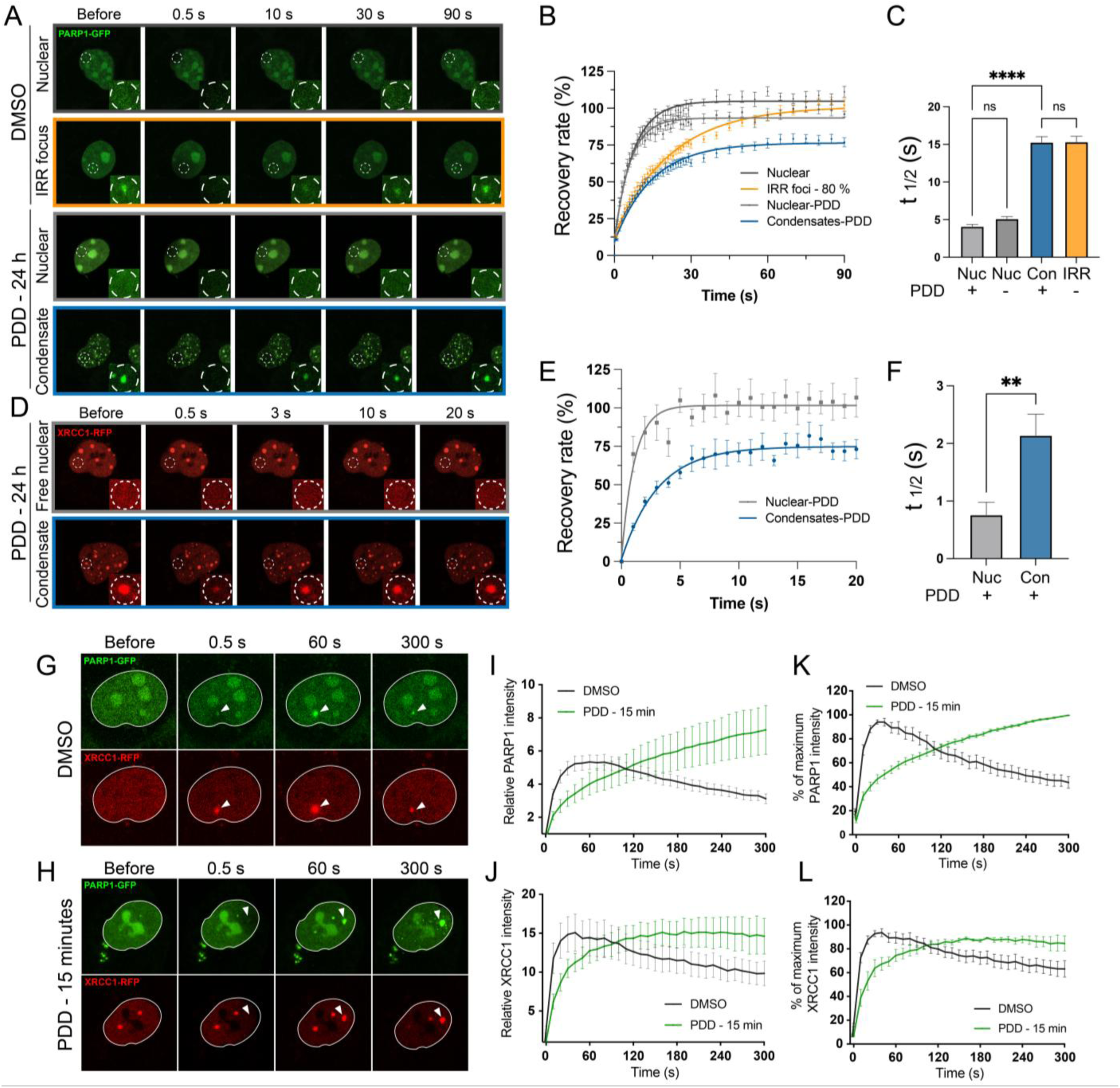
PARGi-induced nuclear condensates are dynamic and resemble DNA damage foci. **A.** Representative images of RPE1WT cells transfected with PARP1-GFP, treated or not with 10µM PDD00017273 for 24 hours, as indicated. White dashed circle highlights the bleached region, lower right square in each image represents a zoom of this region. Images are color-coded corresponding to the representation of the condition in B and C for easier reading. **B.** Calculated FRAP recovery curves for PARP1-GFP from A. Error bars represent SEM. **C.** Half-time (t_1/2_) of the different conditions presented in A. **D.** Representative images of RPE1WT cells transfected with RFP-XRCC1, treated with 10µM PDD00017273 for 24 hours. White dashed circle highlights the bleached region, lower right square in each image represents a zoom of this region. Image is squared with the color corresponding to the representation of the condition in E and F for easier reading. **E.** Calculated FRAP recovery curves for RFP-XRCC1 from D. Error bars represent SEM. **F.** Half-time (t_1/2_) of the different conditions presented in D. **G-H.** Representative images of micro-irradiation-induced PARP1-GFP and RFP-XRCC1 recruitment in PARP1KO RPE1 cells, pretreated approximately 15 minutes with PDD 5µM. White arrowheads point the 405 nm laser damage focus. **I-J.** The relative intensity kinetics of PARP1-GFP recruitment from G and H. Error bars represent SEM. **K-L.** Normalized kinetic curve from I-J with maximum intensity at 100%. Error bars represent SEM. Statistical analysis in C was performed using ordinary one-way ANOVA with Šídák’s multiple comparisons test. Statistical analysis in F was performed using unpaired student t-test. (*p ≤ 0.05, **p ≤ 0.01, ****p ≤ 0.001, ****p ≤ 0.0001).

Next, we directly compared the kinetics of micro-irradiation-induced PARP1 and XRCC1 foci in the presence or absence of PDD. In untreated cells, PARP1-GFP and RFP-XRCC1 rapidly accumulated at damage sites, peaking at ∼40 s, followed by prompt dissociation, consistent with previous reports^46^. In contrast, 15 minutes PDD pre-treatment delayed initial recruitment and markedly impaired foci resolution. PARP1 accumulation was slowed (time to reach relative intensity of 5 extended from ∼40 s to ∼140 s), and XRCC1 accumulation was similarly delayed **(Fig. 3G-J)**. Rather than dissipating, PARP1 foci continued to accumulate over the 300 s observation window, reaching higher intensity (∼8 at 5 min) **(Fig. 3K)**, while XRCC1 foci remained persistently elevated **(Fig. 3L)**. The apparent plateau in XRCC1 signal likely reflects saturation of the measurement. Notably, PARG inhibition also induced pan-nuclear PARP1 condensation at later time points **(Fig. 3G, 300 s)**, consistent with recent observations by Jensen and colleagues^4^. Together, these results indicate that PARG inhibition does not immobilize PARP1 or XRCC1 but instead delays the resolution of PAR-dependent repair assemblies, leading to their persistence and accumulation long after repair completion.

### In PDD-induced condensates, XRCC1 binds to PAR via its BRCT1 domain and recruits LIG3, POLB, and PARP2

Because XRCC1 binds PAR through its BRCT1 domain and itself is also a substrate of PARylation, we next tested whether PARylation of XRCC1 is required for condensate formation. We expressed an RFP-XRCC1^BRCT1^, which lacks PARylation sites in the linker between BRCT1 and BRCT2^62,63^ and the N-terminal POLB-interacting and C-terminal LIG3-interacting domains and assessed its behavior following 24 hours of PDD treatment. RFP-XRCC1^BRCT1^ formed robust nuclear foci that co-localized with PARP1-GFP, comparable to full-length XRCC1 **(Fig. 4A-B versus Fig. 2A-B)**. These results indicate that condensate formation is independent of XRCC1 PARylation and does not require its interactions with POLB via the N-terminal region or LIG3 via the BRCT2 domain. Given gRNAs targeting LIG3 and POLB were also synergistically lethal with PDD, we next asked whether XRCC1 recruits LIG3 and POLB into the condensates. Indeed, both LIG3-GFP and POLB-GFP co-localized with RFP-XRCC1 in PDD-induced pan-nuclear condensates **(Fig. 4C-F**). Consistent with our previous findings that XRCC1 recruits PARP2 to DNA damage foci through a BRCT2-mediated interaction^48^, PARP2 was also enriched in these condensates **(Fig. 4G-H**). Moreover, LIG3-GFP and POLB-GFP formed PDD-induced condensates in cells expressing only endogenous PARP1 and XRCC1 **(Fig. 4I-L)**, and LIG3 condensation was abolished in XRCC1^KO^ cells **(Fig. 4I-J**). Taking together, these results support a model in which prolonged PARG inhibition causes accumulation of PAR-containing condensates that recruit XRCC1 and through XRCC1, co-accumulation of LIG3, POLB, and PARP2 at post-repair sites.

**Figure 4.**
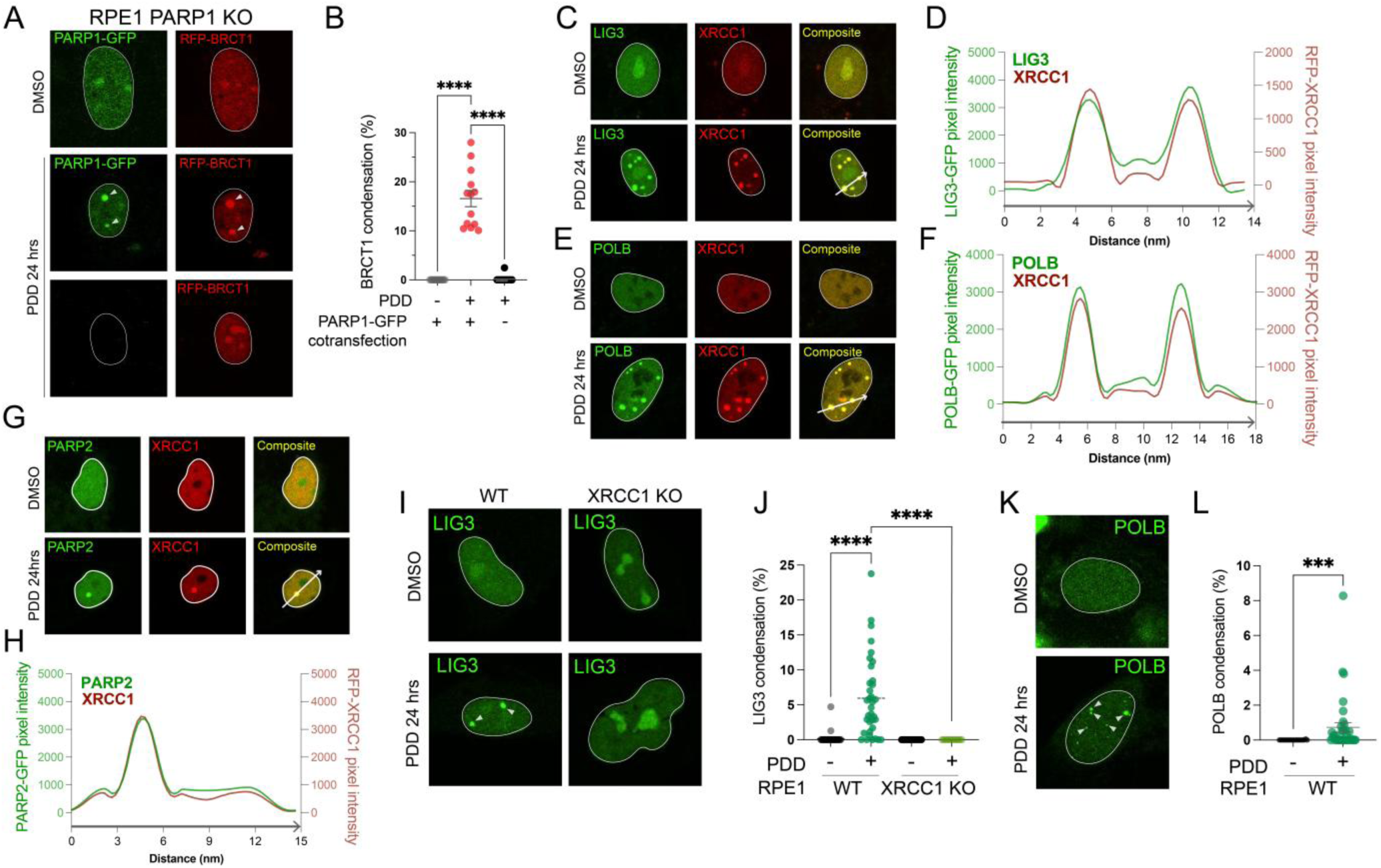
XRCC1 recruits LIG3, POLB, and PARP2 into PARGi-induced nuclear condensates. **A.** Representative images of PARP1-GFP and RFP-XRCC1^BRCTI^ nuclear distribution in DMSO or PDD treated RPE1 PARP1KO cells. Lower panel is representative for RFP-XRCC1^BRCTI^ only transfected cells. White arrows point at the condensates. **B.** Quantification of condensates intensity over the total nuclear intensity. **C-H.** Representative images of nuclear distribution of indicated transfected proteins after 24 hours PDD treatment at 10µM (**C,E,G**). Composite represents green and red merged images, colocalization in yellow. White arrow represents the quantification profile highlighted in right panels **D,F,H** - intensity profiles for indicated proteins. **I.** Representative images of GFP-LIG3 nuclear distribution in DMSO or PDD treated RPE1WT or RPE1 XRCC1-KO cells. White arrowheads point at the condensates. **J.** Quantification of LIG3 relative condensates intensity over the total nuclear intensity. **K.** Representative images of GFP-POLB nuclear distribution in DMSO or PDD treated RPE1WT cells. White arrows point at the condensates. **L.** Quantification of POLB relative condensates intensity over the total nuclear intensity. Statistical analysis in B and J was performed using ordinary one-way ANOVA with Šídák’s multiple comparisons test. Statistical analysis in L was performed using unpaired student t-test. (*p ≤ 0.05, **p ≤ 0.01, ****p ≤ 0.001, ****p ≤ 0.0001).

### PARGi impedes SSBR factors recruitment at *de novo* DNA damage foci, leading to MMS sensitivity

To determine the functional consequences of the PAR-dependent condensates, we assessed micro-irradiation–induced de novo repair foci formation after 4 or 24 hours of PDD treatment, when nuclear condensates are prominent **(Fig. 2)**. Consistent with the sequestration model, prolonged PARG inhibition markedly impaired recruitment of repair factors to DNA damage sites. After 24 hours of PDD treatment, laser-induced PARP1 foci were nearly abolished **(Fig. 5A-C)**, in sharp contrast to the enhanced or sustained accumulation observed following short-term (15 minute) inhibition **(Fig. 3)**. XRCC1 showed a similar trend: peak foci intensity decreased progressively with increasing PDD exposure, from ∼15 in untreated cells to ∼10 after 4 hours and ∼3 after 24 hours **(Fig. 5D**), remaining low throughout the 5-minute observation window. Linear regression analysis revealed a strong inverse correlation between XRCC1 nuclear condensation and IRR-induced XRCC1 recruitment (R=0.998, **Fig. 5E)**, indicating that PARGi-induced condensates functionally sequester PARP1 and, more prominently, XRCC1 and its partners, thereby limiting their availability at newly arising DNA lesions.

**Figure 5.**
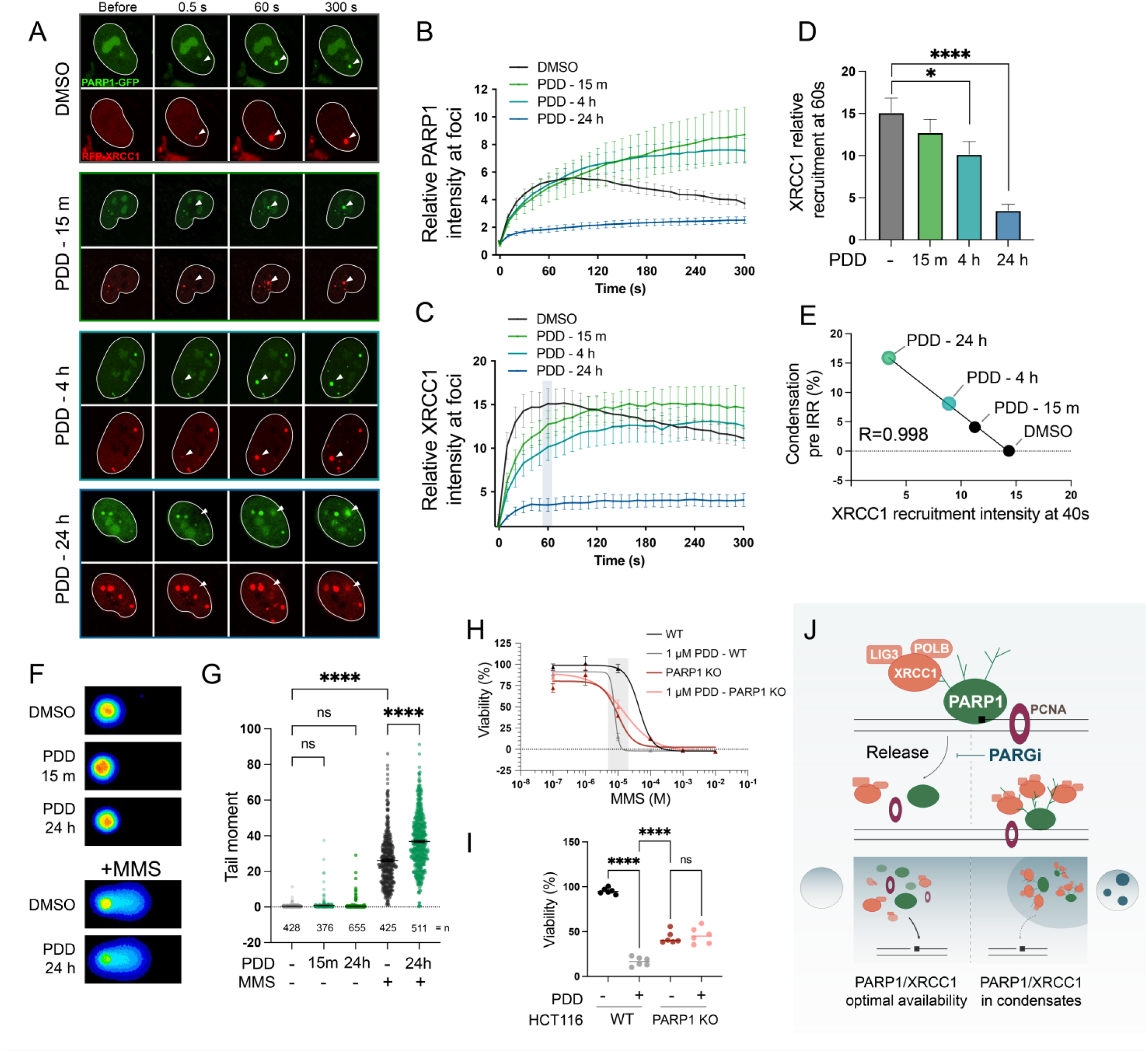
PARGi-induced nuclear condensates sequester XRCC1/PARP1 and delay SSB repair. **A.** Representative images of micro-irradiation-induced PARP1-GFP and RFP-XRCC1 recruitment in PARP1KO RPE1 cells, using indicated PARGi treatment at 5µM (15 minutes, 4 hours) or 10µM (24 hours). White arrowheads point the 405nm laser damage focus. **B.** Relative intensity kinetics of PARP1-GFP recruitment from A. Error bars represent SEM. **C.** Relative intensity kinetics of RFP-XRCC1 recruitment from A. Error bars represent SEM. **D.** Relative intensity of XRCC1 recruitment at 60 seconds post-irradiation from C (grey rectangle). **E.** Simple linear regression representing RFP-XRCC1 relative condensates intensity (from Fig. 2B) over the relative intensity of XRCC1 recruitment at 40 seconds post-irradiation. R represents Pearson correlation score. **F.** Alkaline comet assay representative images for indicated conditions. **G.** Alkaline comet tail moments (= tail length x proportion of DNA in tail) in the indicated conditions. n shows the number of cells for each samples. **H.** Survival of WT and PARP1KO HCT116, relative to untreated condition, in the indicated conditions. **I.** Detailed histogram of H grey rectangle, survival at 10µM MMS. Error bars represent SD. All statistical analysis (D, G, I) were performed using ordinary one-way ANOVA with Šídák’s multiple comparisons test (*p ≤ 0.05, **p ≤ 0.01, ****p ≤ 0.001, ****p ≤ 0.0001) **J.** Model for PARGi mechanisms of action: PARGi initially delays the release of PARP1 at the damage site, induces aberrant PAR chains production and SSBR proteins (in orange) recruitment; PARP1-XRCC1-LIG3-POLB condensates form upon long-term PARGi, disrupting SSBR protein nuclear homeostasis; Further DNA break repair is impaired by less availability of condensates-sequestered proteins.

We next examined whether this sequestration compromises BER. In the absence of exogenous damage, short- or long-term PARG inhibition did not significantly increase alkaline comet tail moments, consistent with the absence of PCNA-positive foci and indicating minimal accumulation of unrepaired breaks **(Fig. 5F-G)** with > 98% of cells having a tail moment below 2, similar to untreated **(Fig. S4A)**. However, upon challenge with the base-damaging agent methyl methanesulfonate (MMS), cells pre-treated with PDD for 24 hours exhibited delayed repair, as evidenced by increased comet tail moments **(Fig. 5F-G)**. Consistently, low dose 1 µM PDD sensitized cells to MMS **(Fig. 5H-I)**, but not the double-strand break (DSB) –inducing agent etoposide **(Fig. S4B),** indicating a selective defect in BER and sequestration of BER, but not DSB repair factors. Importantly, PARP1 loss conferred resistance to PARG inhibition **(Fig. S4C-D)** and abolished the synergistic cytotoxicity between PDD and MMS **(Fig. 5H-I)**, demonstrating that PARP1 activity is required for PARGi-induced MMS sensitivity. Together, these findings indicate that PARG inhibition delays the disassembly of XRCC1–LIG3–POLB repair complexes, leading to their functional sequestration and a selective impairment of single-strand break repair.

## Discussion

PARG inhibitors have emerged as a promising alternative to PARP1/2 inhibitors for modulating NAD⁺-dependent DNA damage responses. However, their mechanisms of action and potential biomarkers for response in proliferating cells remain poorly defined. Using parallel CRISPR screens with multiple PARGi and PARPi, our findings showed that PARP and PARG inhibition have distinct cellular consequences in proliferating cells and characterized a previously unrecognized role for PARG in the timely disassembly and recycling of PAR-binding proteins, particularly XRCC1-LIG3-POLB, after DNA repair. Specifically, we demonstrated that PARG is not required for the initial formation of DNA damage foci and SSB repair, evidenced by the robust initial recruitment after short PDD treatment, the lack of comet tails in PDD-treated cells and the absence of PCNA in the nuclear condensates. Instead, prolonged PARG inhibition delays the disassembly of PARP1-generated, PAR-dependent nuclear condensates following DNA break repair^64,65^. Although the dominant lesions may vary across cell types and conditions, uracil excision by UNG is a major source of endogenous PARP1-activating lesions in our system, as evidenced by the resistance conferred by UNG loss. In addition, the PARGi sensitivity is driven primarily by excessive activation of PARP1, but not PARP2, in the nuclear, as the loss of nuclear NMNAT1, but not cytosolic NMNAT3 or mitochondrial NMNAT2, or the upstream nicotinamide-generating enzyme NAMPT, confers resistance. Persistent condensates sequester PAR-binding proteins, including XRCC1 and its obligatory partners LIG3 and POLB, in a time- and dose- dependent manner. Prolonged PARG inhibition (> 24 hours) depletes the free nuclear XRCC1-LIG3-POLB pool, delaying and attenuating their recruitment to newly induced DNA breaks **(Fig. 5J)**. This model explains the genetic vulnerability of PARG-inhibited cells to the loss of PAR-binding proteins, including XRCC1-LIG3-POLB as well as ALC1, which is distinct from PARP1 inhibition and highlights the critical importance of keeping the PAR-dependent repair foci “transiently”.

While PARP inhibition also delays SSB repair by XRCC1-LIG3, why PARPi preferentially sensitize HR-deficient cells, but PARGi do not? Instead, PARGi selectively sensitize cells with defects in XRCC1, LIG3, and POLB as well as ALC1, while UNG loss confers resistance **(Fig. 1)**. The key difference is whether the breaks were repaired or not and whether PARP1/2 is actively trapped to prevent repair. Unlike canonical micro-irradiation-induced repair foci, PARGi-induced nuclear foci frequently lack PCNA, a PAR-independent and sensitive marker of ongoing DNA strand breaks^66^ **(Fig. 2)**, suggesting the lack of ongoing breaks. Although we cannot exclude the possibility that those PARGi-induced condensates may be initiated by transcription in the absence of *bona fide* strand breaks^64^, UNG deletion confers significant resistance, suggesting that at least a significant fraction might be initiated via uracil processing by UNG. Nevertheless, unlike PARPi, short-term PARG inhibition (15 min and 4 hours) did not markedly impair the initial recruitment of XRCC1 and PARP1 nor cause significant comet tails **(Fig. 5)**. While PARP1 condensates have been described in PARGi-treated cells exposed to exogenous DNA damage^4^, we showed that those condenses accumulate independently of external damage and also contain XRCC1 and its partners **(Fig. 2)**. FRAP analyses further show that PARP1 and XRCC1 within PARGi-induced condensates exhibit dynamic exchange kinetics comparable to *bona fide* micro-irradiation-induced repair foci, rather than irreversible protein aggregates **(Fig. 3)**. Together these observations support a model in which PARG is dispensable for initial strand break repair but is required for the timely disassembly of PAR-based, transient DNA repair centers following successful strand ligation. Accordingly, PARGi-induced PARP1/XRCC1 condensates are not associated with unrepaired DNA breaks, in contrast to the persistent PARP1 foci at unligated DNA breaks observed in XRCC1-deficient or PARPi-treated cells. In those latter settings, unligated single-strand breaks can be converted into replication-associated, often single-ended, DSBs, which are preferentially repaired via HR. By contrast, in PARGi-treated cells, PARP1/XRCC1 condensates persist after repair completion, explaining why HR factors emerge as strong synergistic vulnerabilities in PARPi screens but not in PARGi screens.

Although DNA strand breaks can activate both PARP1 and PARP2, our and others^35^ screens consistently show that loss of PARP1 - but not PARP2 - confers resistance to PARG inhibition. Consistently, PARP1-deficient RPE1 cells, despite retaining endogenous PARP2, display a 4-fold reduction in XRCC1 condensation compared to wild-type cells and fail to form XRCC1 BRCT1 condensates after prolonged PARGi **(Fig. 2 and 4)**. Similar findings have been reported *in vitro*, where purified PARP1 and PARP2 differ markedly in their ability to form phase-separated droplets^67^. Several mechanisms likely contribute, as condensate formation depends on PAR chain length, intermolecular interactions, and protein abundance^67,68^. First, PARP1 accounts for ∼80% of DNA damage-induced PAR activity. Additionally, PARP1 and PARP2 differ in the nature of the PAR chains they produce. Our previous analyses using a catalytically inactive PARP2 model suggest that PARP1 and PARP2 primarily undergo auto-PARylation and cannot efficiently PARylate each other^14^. In PARP1, the catalytic domain is positioned in close proximity to the auto-PARylation loop nested between the BRCT and WGR domains, enabling repetitive PARylation and efficient chain extension, a feature absent in PARP2^2,14,69,70^. Finally, condensate formation requires intermolecular interactions. The three zinc finger domains, specific to PARP1 N-terminus, support protein-protein and protein-PAR interactions^67^. Together with the lower cellular abundance of PARP2, these features explain the unique ability of PARP1 to support PARGi-induced nuclear condensate formation. Furthermore, we found that PARP2 is partially sequestered within PARGi-induced condensates (**Fig. 2**), placing it at the opposite end, as a potential synergistic lethal factor with PARG inhibition. These intrinsic differences between PARP1 and PARP2 in PAR chains generation and *in vitro* condensates formation, combined with the dual role of PARP2 as an XRCC1-dependent PAR-binding factor sequestered by PARGi, likely account for the presence of PARP1, but not PARP2 as a significant hit in PARGi-CRISPR screens.

If not HR deficiency, what drives vulnerability to PARG inhibition? Our model points to PAR-binding factors and basal PARP1 activation as key biomarkers. We documented the sequestration of XRCC1-LIG3-POLB in PAR-dependent nuclear condensates and depletion of their freely available nuclear pool. This sequestration is PARP1-, PAR-, and XRCC1-dependent and is more pronounced for LIG3, which relies entirely on XRCC1 for its nuclear stability, than for POLB. By contrast, phosphorylation-dependent XRCC1 interactors, APLF and PNKP^71^, do not emerge as significant hits, suggesting that both consistency and binding affinity to XRCC1 modulate the degree of sequestration. But XRCC1 and its friends are certainly not the only factor sequestered by PAR. Another PAR-binding protein, ALC1, also emerges as a significant hit, alongside PARG itself, a macrodomain-containing PAR-binding protein, and ARH3, which binds terminal ADP-ribose to catalyze its hydrolysis. These observations suggest that additional PAR-binding factors may be similarly trapped and contribute to cellular sensitivity. While PARG dissolves PAR, the extent and kinetics of PAR accumulation in PARG-inhibited cells are likely modulated by basal nuclear PAR levels. Accordingly, the loss of UNG, PARP1, and NMNAT1 all confer resistance. Together, these findings define a PARGi-specific vulnerability landscape centered on PAR homeostasis and the recycling of PAR-binding proteins, mechanistically distinct from the replication-associated DNA damage and HR dependency characteristic of PARP inhibition.

Finally, while most studies have emphasized the initiation of PAR-dependent DNA damage responses and the recruitment of repair factors, our findings highlight the equally critical importance of foci disassembly and repair factor recycling. This provokes a fundamental question: what is the biological role of the transient assembly of PAR-dependent nuclear foci? PAR chains are highly negatively charged and attract many nucleic acid proteins. In addition to repair factors, multiple RNA-binding proteins^68,72^, and up to 70% of transcription factors containing nucleic acid-binding domains (e.g., zinc finger, helix–loop–helix) tested can localize to PAR-dependent foci after radiation^73^. Although NAD+-dependent ADP-ribosyltransferases are evolutionarily ancient, DNA break-activated PARPs are absent from prokaryotes and simple eukaryotes such as yeast; appearing as PARP1 orthologs in plants and insects; and further expanded to include PARP2 in vertebrates with increasing genome size and nuclear complexity. From an evolutionary perspective, we speculate that PARP1-driven, PAR-dependent nucleic acid-protein assemblies function as mobile repair/transcription hubs, effectively recreating high local concentrations of nucleic acid-binding factors within the much larger and spatially complex mammalian nucleus. Transient concentration of DNA repair, RNA processing, and transcription factors within PAR-rich condensates enables efficient lesion recognition and processing without requiring an increase in total nucleic-acid binding proteins abundance. In this framework, however, timely disassembly of PAR-dependent foci is as essential as their formation, as it restores the freely diffusible nuclear pool and preserves genome-wide responsiveness. Our findings suggest that disruption of this balance via PARG inhibition uncouples assembly from disassembly, leading to pathological factor sequestration and cellular vulnerability. In addition to XRCC1 and its partners examined in detail here, PARG inhibition might sequester other PAR-binding proteins beyond ALC1, PARG, and ARH3. Indeed, the top synergistic lethal hits in our screen also include several DNA repair factors that are known to form PAR-dependent foci (e.g., RFC, ZC3H8, and RAD17)^73^ as well as RNA-binding proteins (e.g., DHX37 and RBM10) **(Table S1)**. Notably, some PAR-binding proteins that are themselves essential (e.g., DDX21 and ribosomal proteins) may be underrepresented in the screen, despite being sequestered upon PARG inhibition^73^ .Together, these findings raise the possibility that PARG inhibition induces widespread sequestration of PAR-binding proteins, extending beyond canonical DNA repair factors. Such broad functional depletion may explain why PARG is essential, whereas PARP1 is dispensable. Future studies would be necessary to determine which of them might represent candidate predictive markers for PARG inhibition.

## Acknowledgments

We thank Drs. Li Lan, Keith Caldecott, and John A. Tainer for sharing critical reagents, cell lines, and chemicals for the studies. We greatly appreciate discussions and suggestions from Drs. Junjie Chen, John Pascal, Hanwen Zhang, and the other members of the Zha lab. The work is supported by NIH CA226852, CA271595, CA174653, CA293675, and CA275184 to SZ; NIH P30CA013696 to HICCC.

## Author contributions

I.D. and S.Z. conceived the project and wrote the manuscript with input from other co-authors. B.L carried out the CRISPR screens, Y.W. performed the initial analyses of the CRISPR Screen. X.L. generated the XRCC1-BRCT1-RFP and the RFP tagged-PARP2 plasmids. I. D. carried out all the quantitative live cell imaging studies and the sensitivity studies presented in the manuscript with assistance from C. Z.

## Material and methods

### Cell lines and plasmids

RPE1 and immortalized MEFs were cultured in DMEM medium (GIBCO, Cat. 12430062), HCT116 were cultured in McCoy’s 5a Medium Modified (ATCC 30-2007), both supplemented with 15% fetal bovine serum (Hyclone, SH30071.03), MEM non-essential amino acids (Gibco, 11140-050), 1 mM sodium pyruvate (Gibco, 11360-070), 2 mM L-glutamine (Gibco, 25030-081), 120 μM β-mercaptoethanol (Fisher, 03446I-100) and 50 U/ml penicillin/streptomycin (Gibco, 15140122). RPE-1 PARP1-KO, PARP2-KO and immortalized MEFs were generated in Shan Zha’s lab. RPE-1 XRCC1-KO is a kind gift from Dr. Keith Caldecott’s lab at the University of Sussex. HCT116 PARP1-KO was generated as detailed in the generation of HCT116 PARP1 knockout clones methods section. PARP1 GFP-tagged plasmid was described previously^46^. PARP2 GFP-tagged plasmid was described previously^61^. RFP-tagged XRCC1, EGFP-tagged LIG3, and POLB were kind gifts from Dr. Li Lan at Duke University.

### CRISPR screen

CRISPR screen workflow was described in Menolfi *et al*., 2023^38^. Briefly, EµBcl2+ v-abl kinase-immortalized murine B cell line was infected with lentivirus encoding doxycycline-inducible SpCas9 (Addgene Plasmid #50661). A single clone was selected for high SpCas9 inducibility after doxycycline treatment (Sigma Aldrich, D9891, 3 µg/ml for 3 days) and expanded. 100 million cells were infected with a BFP-expressing gRNA lentiviral library (Addgene Pooled Library #67988). After expanding for 4 days, at least 35 million BFP+ cells were sorted to ensure 250–300x library coverage. The sorted cells were expanded and DNA from 3 replicates (30 Million cells each) was collected: *before doxycycline* controls. The rest was treated with 3 µg/mL doxycycline for 5 days, and DNA from 3 replicates was collected: *after doxycycline* controls. The rest was divided into 30 million cells plates for triplicate 6-day treatment with IC90 olaparib (0.69 µM), niraparib (0.27 µM), talazoparib (2.58 nM), PDD00017273 (43.62 µM), JA2131 (25.8 µM), COH34 (9.72 µM, screen 1; 11.73 µM, screen 2) and DMSO before DNA collection. gRNAs from all collected samples were amplified before sequencing as described by Konermann *et al.*, 2017^74^. Amplification was performed using primers with Illumina sequencing adapters and unique barcodes for each sample. The libraries were pooled and sequenced on a NextSeq 550 machine (Illumina), and results were analyzed via the MAGeCK and MAGeCK FluteMLE pipeline as detailed in the technical support page (https://sourceforge.net/p/mageck/wiki/Home/) using the DMSO samples as the baseline. Heatmaps were generated on Prism GraphPad 10 and modified on Adobe Illustrator 2025.

### MTT assay

For viability assays, cells were harvested, counted, plated in 96-well plates (400 cells per well) and treated 6 hours after plating. All conditions were done at least in triplicate. After 5 days, MTT reagents from Roche-Sigme Aldrich Cell proliferation kit I (Cat: 11465007001) were added following the manufacturer’s recommendations. After overnight incubation, reading of the 560nm absorbance of each well was performed using Promega Plate Reader GloMax. Media-only blank measures were subtracted from the absolute absorbance reads before normalization: treated divided by the average of three untreated conditions. When needed, IC90 was determined using the function “log(inhibitor) vs. normalized response - variable slope” equation within nonlinear regression in Prism software package.

### Live cell imaging for condensation assays, laser micro-irradiation and FRAP

RPE1 cells or MEFs were plated at 200 000 cells per µ-dish 35 mm glass-bottom plate (IBIDI, cat. 8156) 48 hours before imaging. The day after plating, plasmids encoding GFP- or RFP-tagged PARP1, XRCC1, POLB, LIG3, PARP2, or PCNA were transfected using Lipofectamine 2000 (Invitrogen, cat. 11668019), following manufacturer’s instructions. Cells were cultured in transfection media for around 6 hours before media change for treatment. Nikon Ti Eclipse inverted microscope (Nikon Inc, Tokyo, Japan) was used for imaging, equipped with A1 RMP (Nikon Inc.) confocal microscope system (Nikon Inc.) and Lu-N3 Laser Units (Nikon Inc.). Laser micro-irradiation, photo-bleaching for FRAP, and time-lapse imaging were conducted via the NIS Element High Content Analysis software (Nikon Inc.). Laser micro-irradiation was performed using 405 nm laser. PARP1-GFP photobleaching experiments were performed using 488 nm laser. RFP-XRCC1 photobleaching experiments were performed using 561 nm laser. If photobleached zone was previously irradiated, photobleaching was performed 60 sec after laser micro-irradiation.

### Condensates quantification

A custom macro in Fiji (ImageJ) was developed to quantify the condensates per nucleus while accounting for both condensate size and intensity. Images were first subjected to automatic thresholding using the “Default dark, no-reset” algorithm. To determine nuclear background intensity, a pixel intensity < 70 ∼200 (16-bit scale) and size ≥100 pixels were applied. To be considered as condensates, the particle has to be ≥0.01 pixels and with pixel intensity > 700–2000 (∼10-fold brighter than respective nuclear thresholds). Both nuclear (background) and condensate measurements were obtained using the ‘Analyze Particles’ function. For each nucleus, condensate abundance was quantified using integrated density (IntDen): the sum of all condensates within a nucleus, and the “PROTEIN condensate (%)” was calculated as: condensates (IntDen) / nucleus (IntDen). For colocalization analyses, a straight line was drawn on Fiji (ImageJ) and “plot profile” function was used to generate the profile intensity for each protein. Curve was then smoothed using Prism GraphPad 10.

### Micro irradiation recruitment kinetics and FRAP quantification

To quantify the dynamics of protein recruitment and recover after photobleaching in specific regions of interest (ROIs) a custom Fiji (ImageJ) macro was used. All time-lapse images were first aligned using the StackReg plugin (Rigid Body transformation) to correct for potential drift. Then, maximal intensity projection was used to manually define ROIs: the nucleus, the outside of the nucleus (background) and the damaged/photobleached spot. The macro recorded the integrated density for each ROIs (nucleus, background, spot, and full image) across all time points. To quantify the recruitment kinetics, background signal was subtracted from all measurements. Then, intensities were normalized to their initial values. Kinetics of recruitment was calculated as spot intensity/nuclear intensity-spot intensity (excluding the spot), providing a measure of protein enrichment at damage sites. For FRAP. All data were normalized on pre-bleached intensity. Non-linear fit and one-phase association Prism GraphPad 10 functions were used to determine t_1/2_.

### Comet assay

The alkaline comet assay was performed using the Trevigen CometAssay Kit (Cat: 4250-050-K). Following treatment, cells were trypsinized, resuspended in culture medium and counted. A total of 100,000 cells per sample were pelleted by centrifugation, washed twice with ice-cold PBS (Ca²⁺/Mg²⁺-free), and resuspended in 50 µL of cold PBS. Low-melting-point agarose (LMAgarose) was melted in boiling water for 5 min before being maintained at 37°C. Cells were mixed with molten LMAgarose (1:9 ratio) and immediately transferred onto CometSlides (50 µL per well). Slides were placed in a humidified chamber and incubated at 4°C in the dark for 30 min. Lysis was performed overnight at 4°C in pre-chilled lysis solution (Trevigen). Then, slides were immersed in freshly prepared alkaline unwinding solution (200 mM NaOH, 1 mM EDTA, pH >13) for 1 hour at 4°C in the dark. Electrophoresis was conducted in alkaline electrophoresis solution (200 mM NaOH, 1 mM EDTA, pH >13) at 21 V / 280-300 mA for 30 minutes. After electrophoresis, slides were rinsed twice in distilled water for 5 min each, dehydrated in 70% ethanol for 5 min, and dried at 37°C for 30 min. Before imaging, slides were stained with ethidium bromide (20 µg/mL, 10 µL per slide). Comets were visualized using a Nikon Eclipse fluorescence microscope with 10x lens. DNA damage was quantified using image analysis software Comet Score 2.0 (http://rexhoover.com/index.php?id=cometscore), measuring tail moment.

### Western Blotting

After harvesting the cells, lysis was performed in a buffer containing 50 mM HEPES (pH 7.5), 250 mM NaCl, 5 mM EDTA, 1% NP40, 1 mM DTT, 1 mM PMSF and EDTA-free Protease Inhibitor Cocktail (Roche). After 30 minutes incubation at 4°C, the total cell lysate was obtained by taking the supernatant from a centrifugation (10,000 xg, 15min, 4°C). A 10% SDS-PAGE gel was used for migration (150V, 1 hour, room temperature) before transfer on PVDF membrane (30V, overnight. 4°C). 5% milk in TBS-Tween buffer was used for blocking and primary antibody. ThermoFisher Pierce™ ECL Western Blotting Substrate was used for revelation and BioRad chemidoc was used for imaging. PARP1 antibody from Cell Signaling (9542, 1:1000) detected the protein. Trichloro Ethanol was added to the gel for stain-free revelation.

### Generation of HCT116 PARP1 knockout clones

PARP1 knockout clones were generated using IDT Alt-R CRISPR-Cas9. Two independent alt-R crRNAs targeting PARP1 (5’-GAG TCG AG TAC GCC AAG AGC GGG-3’, and 5’-ATT GAC CGC TGG TACC ATC CAG G-3’) were annealed with tracrRNA-ATTO 550 by warming up samples at 95°C for 5 minutes, then cooling to room temperature for 30 minutes to form gRNAs. gRNAs were incubated with Alt-R Cas9 Nuclease V3 at room temperature for 15 minutes to form RNP. HCT116 cells were harvested, washed with PBS, and resuspended at a density of 200.000 cells. Cells were pelleted and resuspended in 20 µL of Nucleofector Solution (SF Cell Line, Lonza) supplemented with both RNP complexes. Electroporation was performed using the Nucleofector 4D device (program EN-113). Transfected cells were immediately transferred to prewarmed culture media in a 12-well plate and incubated at 37°C with 5% CO2. After 48 hours, cells were seeded at 3 different limiting dilutions in 96-well plates to isolate single-cell clones. After 12-14 days, single clones were expanded before lysis and Western Blot to confirm PARP1 depletion.

## Supplementary Table and Figure Legends

**Supplementary Table 1. CRISPR screen results.**

After analysis by MAGeCK-Flute pipeline, the following metrics are reported for each gene: Beta-score, Z-score, p-value, false discovery rate (fdr), wald z-score, wald p-value and wald fdr, and for all three PARP inhibitors (talazoparib/BMN673, olaparib, niraparib) and three PARG inhibitors (PDD00017273, JA2131, and COH34 (2 independent screens)).

**Supplementary Figure 1.**
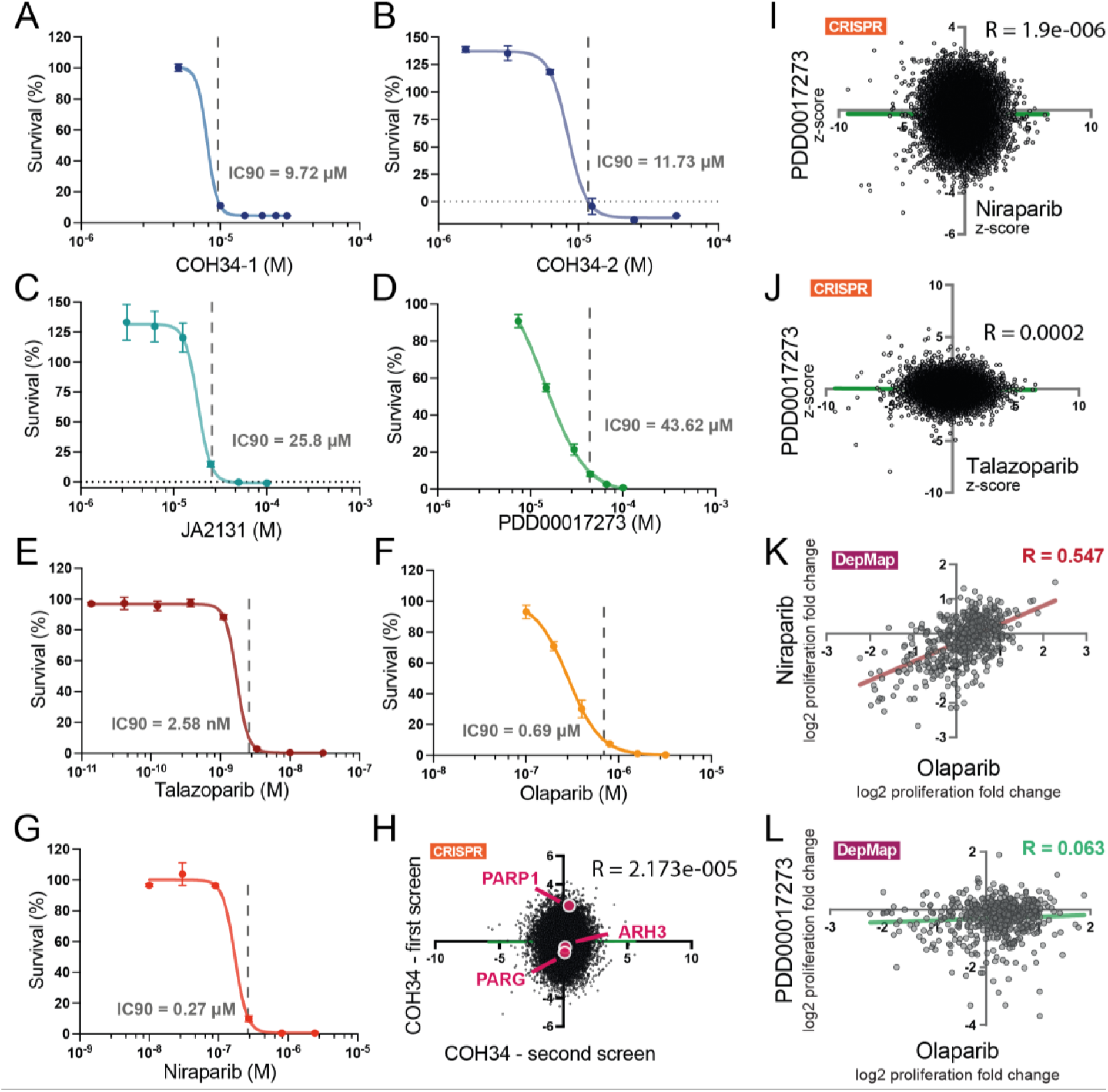
Data supports the CRISPR screen and validation. A-D. MTT proliferation assay of v-abl cells treated with olaparib (A), niraparib (B), talazoparib (C) and PDD00017273 to determine IC90 of the drugs (dashed grey line). **E-G.** Scatter plots of total CRISPR genes Z-scores for COH34 - replicate 1 vs COH34 - replicate 2 (E), PDD vs niraparib (F), PDD vs talazoparib (G) IC90 treatments. Simple linear regression and Pearson correlation test were performed to determine R. **H-I.** Scatter plots of DepMap sensitivity scores (PRISM repurposing data) of 536 cell lines to the indicated compound. Simple linear regression and Pearson correlation test were performed to determine R.

**Supplementary Figure 2.**
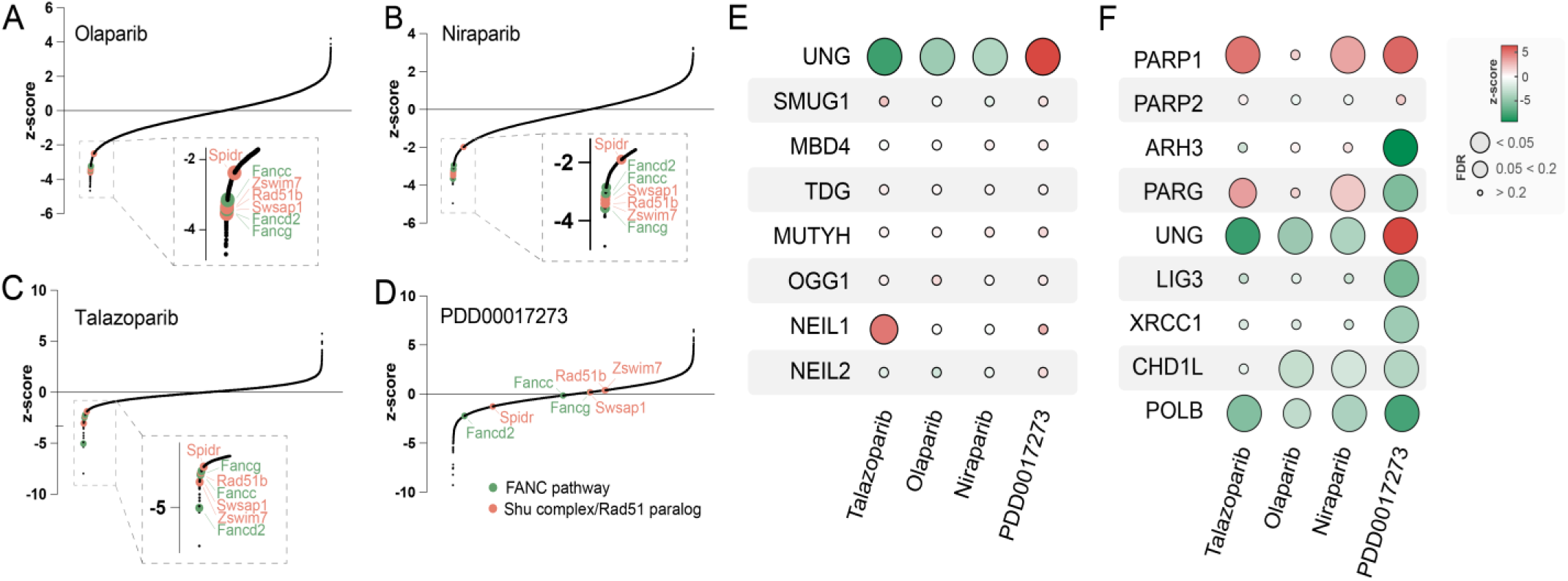
Additional CRISPR screen results. A-D. CRISPR screen z-scores ranking of HR related co-essential genes with olaparib (A), niraparib (B), talazoparib (C) and PDD00017273 (D) treatment. Shu complex or RAD51B related genes are shown in orange, FA related genes are shown in green. **E-F.** Heatmap representing z-score (color range) and False Discovery Rate FDR (size) for the genes targeting indicated in rows, in cells challenged with the inhibitor indicated above, in columns. Panel E. displays glycosylases, and panel F. displays several genes of the SSBR pathway.

**Supplementary Figure 3.**
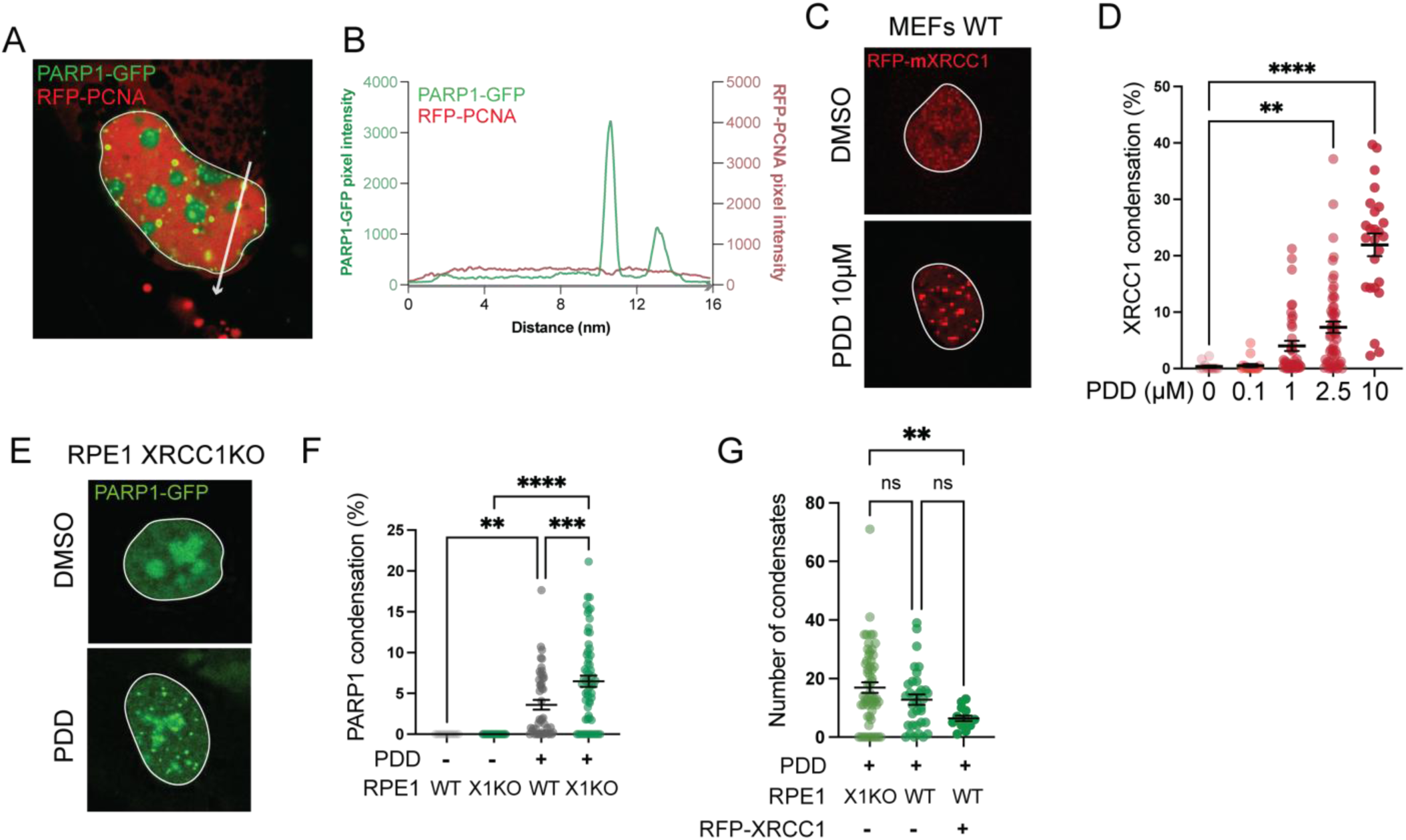
Additional Cell Biology Data support the sequestration model. **A.** RPE1 PARP1KO cells transfected with PARP1-GFP and RFP-PCNA, treated with PARGi for 24 hours (10 µM). White arrow represents the quantification profile highlighted in B. **B.** Representation of intensity profile for PARP1 (green) and PCNA (red). **C.** Representative images of wild type Mouse Embryonic Fibroblasts (MEFs) transfected with RFP-mXRCC1, treated for 24 hours PDD 10 µM. **D.** Quantification of RFP-mXRCC1 condensates intensity over the total nuclear intensity in a concentration-course experiment at 24 hours PDD treatment. **E.** Representative images of PARP1-GFP nuclear distribution in DMSO or PDD treated RPE1 XRCC1-KO cells. **L.** Representation of the condensate intensity over the total nuclear intensity of the indicated protein. Gray data were already used in Figure 2J. **G.** Count of the number of condensates per nucleus in the indicated cell lines and treatment conditions.

**Supplementary Figure 4.**
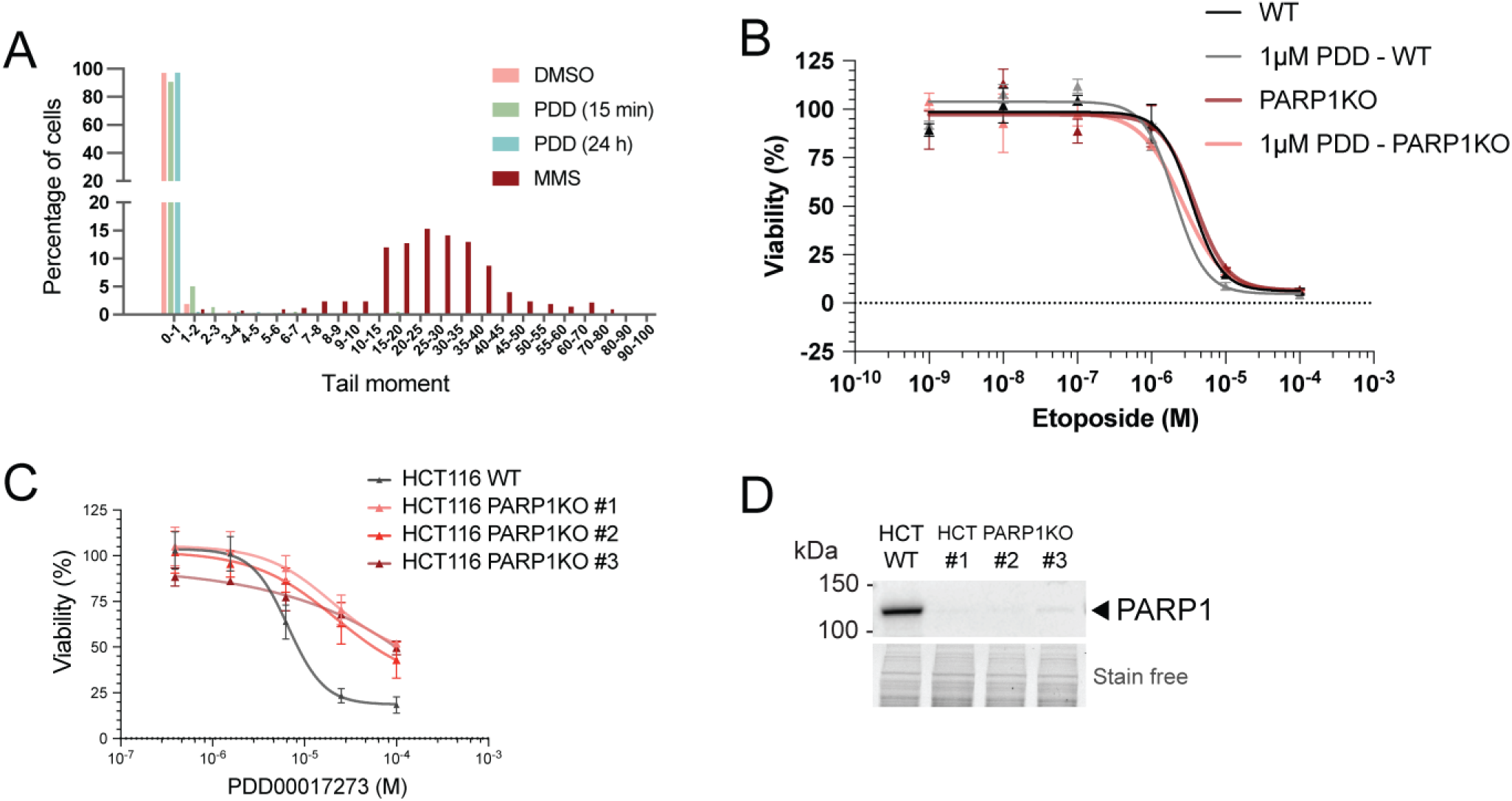
Data supports the selective SSB repair defects in PDD-treated cells. **A.** Representation of Comet assay tail moments (from representative images shown in Fig. 5F) with the percentage of cells (Y axis) over the tail moment ranges. As an example, “0-1” in X axis means that cells in this category have a tail moment between 0.00 and 1.00. For the following, “1-2” corresponds to cells that have a tail moment between 1.01 and 2.00. **B.** Survival of WT and PARP1KO HCT116, relative to untreated conditions, in the indicated conditions. **C.** Survival of WT and PARP1-KO HCT116 clones, relative to the untreated condition, in the indicated conditions. **D.** Western-blot for PARP1-KO generated clones used in C. Gel total proteins (stain-free) are used as the loading control.

